# Testicular gene expression patterns suggest a heterochronic shift underlying *vgll3*-mediated maturation age variation in Atlantic salmon

**DOI:** 10.1101/2025.03.05.641643

**Authors:** Ehsan Pashay Ahi, Jukka-Pekka Verta, Johanna Kurko, Annukka Ruokolainen, Paul Vincent Debes, Craig R. Primmer

## Abstract

Heterochrony, or shifts in developmental timing, drives phenotypic diversity within and between species and shapes life history traits that can be selected for in changing environments which in turn promotes population resilience. Despite its importance, the molecular basis of heterochrony remains largely unknown. Mutations in “heterochronic genes” that regulate these processes can induce stable timing shifts, impacting important life history traits like pubertal timing. Heterochronic shifts in gene transcription are often tissue-specific and in mammals, for example, the testis shows the most pronounced heterochrony across species, especially during spermatogenesis. Age at maturity is a key adaptive trait across species, with *vgll3*, a Hippo pathway co-factor, as a main determinant in Atlantic salmon. The roles of *vgll3* in maturation timing, adiposity, and energy storage are evolutionarily conserved across fish and mammals. Recent studies in salmon show *vgll3* alleles; *early* (E) and *late* (L), affect reproductive gene expression, reinforcing its role in regulating developmental timing. This study examines whether *vgll3* influences testicular heterochrony in Atlantic salmon by analyzing Hippo pathway-related gene expression in E and L genotypes. We observed heterochronic divergence in Hippo pathway gene transcription, indicating accelerated spermatogenesis-linked changes in the testes of *vgll3*EE* individuals. Since maturation in Atlantic salmon is closely tied to environmental changes, and the Hippo pathway acts as an environmental sensor, these findings suggest that Hippo-*vgll3* shifts may also respond to environmental signals. This positions *vgll3* as a heterochronic gene which is potentially affected by environmental changes (heterokairic), making it an ideal target for studying ecological adaptation linked to heterochrony.

## Introduction

Changes in the timing of developmental events, known as heterochrony, contribute to the phenotypic differences observed within and between species (Rougvie, 2001; Garcia-Ojalvo & Bulut-Karslioglu, 2023). Heterochrony involves shifts in the onset, completion, or rate of developmental events relative to an ancestor. Understanding these timing mechanisms is essential for grasping their evolutionary impact (Smith, 2003). Throughout an individual’s life, environmental stressors may also influence these heterochronic events, a process known as heterokairy, adding another layer of phenotypic diversity at the individual level (Rundle and Spicer, 2016). This becomes especially likely if the molecular signals driving heterochronic events are also responsive to environmental changes (Dubansky, 2018; Spicer et al., 2018). Heterochronic processes give rise to life history trait variation, and may sometimes be the targets of selection even contributing to major evolutionary processes like vertebrate evolution (McNamara, 2012). Alternative life history strategies arising from various traits significantly enhance population diversity, supporting adaptive capacity and resilience (Brommer, 2000). Mutations in genes regulating heterochronic processes, so called “*heterochronic genes”*, if advantageous, could drive stable shifts within populations, potentially becoming key sources of diversity (Rougvie, 2001). A classic example of heterochronic genes is a highly conserved family of RING finger/B-box proteins known as *Lin* genes (Ambros & Horvitz, 1984). These genes regulate the timing of developmental events across various organisms such as pubertal timing (age at maturity) (Rougvie, 2001).

Age at maturity is a key adaptive life history trait in many organisms, but in humans, shifts in maturation timing—leading to early or delayed puberty—can introduce various physical and psychological challenges (Hartge, 2009; Day et al., 2017; Leka-Emiri et al., 2017). One well-studied example of a heterochronic gene affecting age at maturity is the *Lin28a/b* gene, a highly conserved factor influencing body size, pubertal timing, and metabolism from worms to humans (Ong et al., 2009; Zhu et al., 2010). The impact of *Lin28a/b* on pubertal timing appears sex-specific, primarily affecting males, and is linked to metabolic changes related to adiposity (Zhu et al., 2010; Faas et al., 2013; Ouchi et al., 2014; Corre et al., 2016). A recent discovery has shown that *Lin28a/b* can act partly independently of *let-7* by directly inhibiting the Hippo pathway, which is involved in sexual maturation and organ size control, through the induction of *YAP1* expression, a negative regulator of the pathway (Luo et al., 2021; Zou et al., 2022). This finding could explain the role of *Lin28a/b* in all the processes mentioned earlier, as the Hippo pathway is a key regulator of body size (Pan, 2007), adipogenesis (Ardestani et al., 2018; Shen et al., 2022), and sexual maturation via the hypothalamus-pituitary-gonadal (HPG) axis (Sen Sharma et al., 2019; Lalonde-Larue et al., 2022).

In Atlantic salmon, the gene *vestigial-like family member 3* (*vgll3*) is a primary genetic determinant of maturation timing, accounting for over 39% of the variation in age at maturity within natural populations (Barson et al., 2015; Czorlich et al., 2018). Although *vgll3* has not been categorized as a heterochronic gene, its function in salmon physiology closely mirrors the roles of *Lin28a/b*, showing sex-specific effects on age at maturity (Barson et al., 2015; Czorlich et al., 2018; Miettinen et al., 2024), influencing adiposity and seasonal energy storage (House et al., 2023; Ahi et al., 2024), and regulating the Hippo pathway by inhibiting YAP1 (Hori et al., 2020; Kurko et al., 2020) (Fig. 1). Interestingly, like *Lin28a/b*, these functions of *vgll3* appear to be evolutionarily conserved. For example, *vgll3* has been linked to age at maturity in humans (Perry et al., 2014; Cousminer et al., 2016), regulation of adipogenesis in mice (Halperin et al., 2013), sex-specific functions in humans, e.g. sex-biased pathogenesis of autoimmune diseases (Liang et al., 2016). Our recent investigations involving male Atlantic salmon have revealed strong correlations between *vgll3* alleles E and L linked to *early* and *late* maturation, respectively, and the expression patterns of important reproductive genes along the HPG axis (Ahi et al., 2022, 2023; Verta et al., 2024).

**Figure 1.**
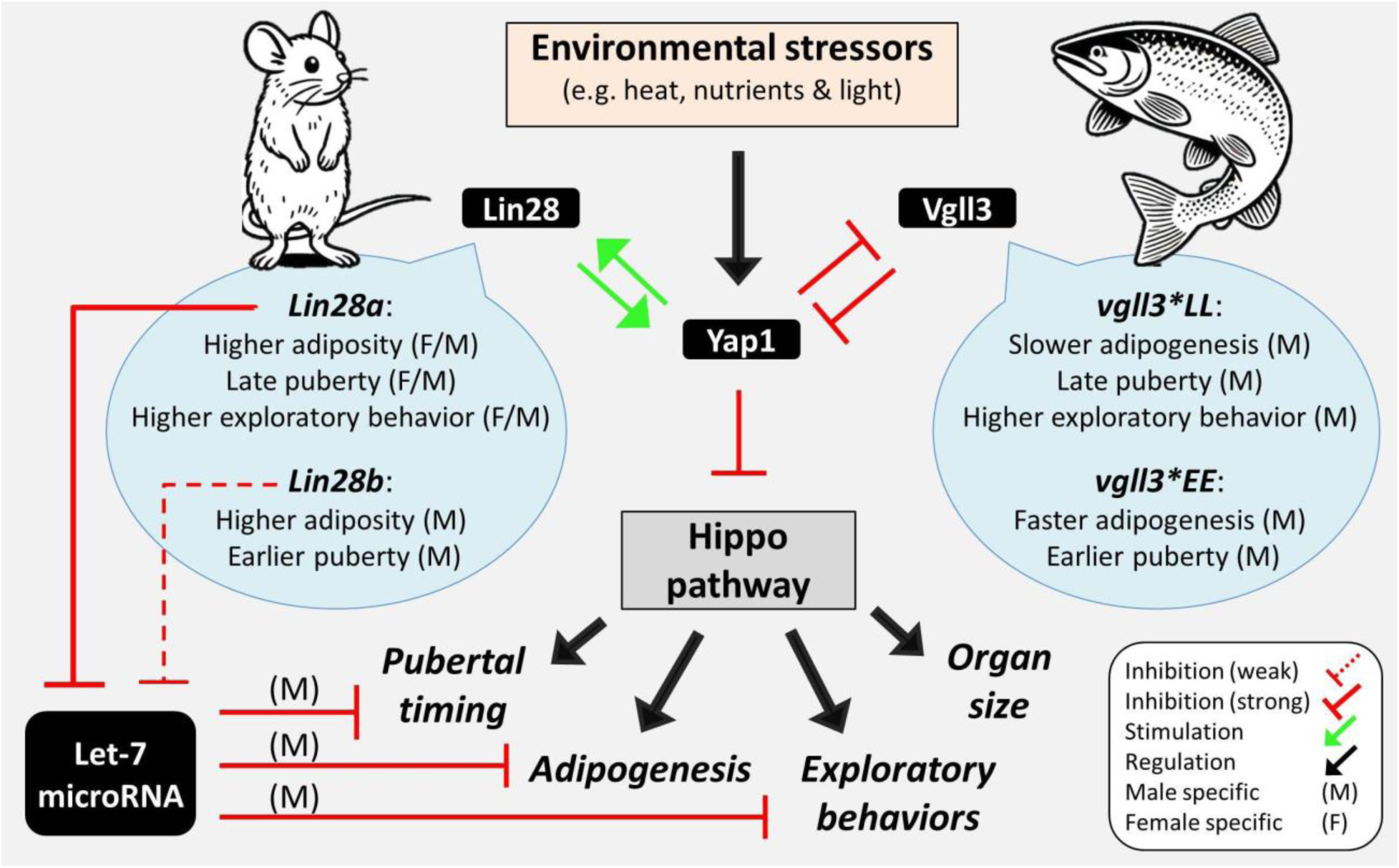
Functional similarities between *Lin28* in mice, and *vgll3* in Atlantic salmon. *Lin28* is a well-known heterochronic gene which regulates pubertal timing across various species and, in mammals, is also associated with adiposity and behavioral effects (i.e. exploratory behavior and boldness). The functional similarities between *Lin28* and *vgll3* in these processes are thought to occur through differential activation of the Hippo pathway (via Yap1 regulation). However, the sex specificity of *Lin28* in mammals appears distinct from that of *vgll3* in salmon, potentially due to *Lin28a/b*’s additional role in regulating the biogenesis of *let-7* (a highly conserved microRNA playing role in developmental heterochrony). *Let-7* influences pubertal timing, adipogenesis, and behavior in a sex-specific manner, with effects more pronounced in males. Since *Lin28a/b* is a stronger suppressor of *let-7*, its induction could override *let-7*’s sex-specific effects in these processes. It is currently unknown whether the male specific effects of *vgll3* is Hippo pathway-dependent or through other molecular players such as *let-7*. However, these similarities indicate a potential role of *vgll3* in developmental heterochrony in Atlantic salmon.

Heterochronic shifts in transcription are not uniformly reflected across all organs (Garcia-Ojalvo & Bulut-Karslioglu, 2023). In mammals, for example, the brain exhibits minimal heterochronic divergence in developmental events, while the testis shows the most pronounced heterochrony across species (Cardoso-Moreira et al., 2019). Particularly, within the testis, transcriptional changes associated with different stages of spermatogenesis display the most significant heterochronic divergence between mammalian species (Cardoso-Moreira et al., 2019). Recent studies have examined the potential functions of *vgll3* in the testis of Atlantic salmon, ranging from isoform-specific (Verta et al., 2020) and cell-specific regulatory roles (Kjærner-Semb et al., 2018) to broader interactions with other pathways involved in testicular development, such as the TGF-β and Wnt signaling pathways (Verta et al., 2024). Beyond these, further pathway-specific molecular investigations would provide deeper insights into the role of the Hippo pathway in testicular maturation. In this study, we aimed to investigate whether the effects of *vgll3* on testis involve heterochronic shifts in the transcription of its downstream molecular targets, i.e. known components of the Hippo pathway as well as their associated interacting partners and signaling molecules. We assessed the testicular expression of these genes in the immature and mature Atlantic salmon, with distinct *early* (E) or *late* (L) maturing *vgll3* genotypes. Through a comparative analysis of maturity status and *vgll3* genotype, we employed methodologies centered on pathways and regulatory networks. The results of our study could be ecologically significant, as the Hippo pathway is emerging as a key sensor of environmental changes, such as temperature and food availability (Shu et al., 2019; Luo et al., 2020). The identification of Hippo-*vgll3*-mediated heterochronic shifts suggests that these shifts may also be influenced by environmental stressors, potentially leading to variation in the timing of related developmental events within an individual’s life—a phenomenon known as heterokairy (Rundle and Spicer, 2016). This finding positions *vgll3* not only as a conserved heterochronic gene but also as a potential heterokairic gene, relevant for studies of ecological adaptation linked to heterochrony within and between populations.

## Materials and methods

### Fish material and tissue sampling

The individuals involved in this study were sourced from the same population (Oulujoki) and cohort as previously detailed in Verta et al. (2020), which provided access to individually PIT-tagged individuals with known *vgll3* genotypes (refer to Verta et al., (2020) for more information on breeding and rearing). For the purposes of this research, male individuals were collected at the age range of 1.5 to 2 years post-fertilization (Debes et al., 2020). The euthanization process involved the administration of an overdose of MS222, after which the entire testes from each male were collected at three different time points, representing distinct stages of maturation development (as described below). Subsequently, these tissue samples were rapidly frozen with liquid nitrogen and stored at a temperature of −80°C until further processing.

*Immature 1*-stage individuals were gathered during the late spring period (5-21 May) and exhibited an average mass of 17.5 g, ranging from 10.2 to 27.4 grams. Their average length measured 12.1 cm, with individual lengths falling within the range of 9.1 to 13.6 cm. These individuals did not display any noticeable signs of testicular maturation, as indicated by a gonadosomatic index (GSI) value of 0.

*Immature 2*-stage individuals were sampled in the summer season (4-17 July). They had an average mass of 33.7 grams, ranging from 21.1 to 76.6 grams, and an average length of 17.3 cm, falling within the range of 14 to 18.5 cm. Some of these individuals exhibited initial indications of phenotypic maturation processes, as reflected by GSI values ranging from 0.0 to 0.3.

*Mature*-stage individuals were collected close to the anticipated spawning period in early autumn (1-15 October). These individuals had an average mass of 82.7 grams, with a range of 60.3 to 120.6 grams, and an average length of 20.8 cm, ranging from 16.4 to 22.7 cm. All individuals in this group displayed well-developed testes, indicative of maturity, with a GSI exceeding 3.

### RNA extraction

RNA was extracted from 24 testis samples using a NucleoSpin RNA kit (Macherey-Nagel GmbH & Co. KG). The collected samples were processed by placing them in tubes containing 1.4 mm ceramic beads from Omni International, along with Buffer RA1 and DDT (350 µl of RA1 and 3.5 µl of 1M DDT). Homogenization was carried out using the Bead Ruptor Elite (Omni International) at a frequency of 30 Hz for a total of 2 minutes, involving 6 cycles of 20 seconds each. The subsequent RNA extraction procedures followed the manufacturer’s guidelines, including an integrated DNase stage to eliminate any remaining genomic DNA. After completion of the process, RNA from each individual sample was eluted using 50 µl of nuclease-free water. RNA quantity was assessed using the NanoDrop ND-1000 instrument (Thermo Scientific, Wilmington, DE, USA), and RNA quality was evaluated with the 2100 BioAnalyzer system (Agilent Technologies, Santa Clara, CA, USA). All samples had a RNA integrity number (RIN) exceeding 7, and 100 ng of extracted RNA from each isolation was used for the hybridization step in the Nanostring panel.

### Nanostring nCounter panel

The NanoString nCounter technology is a versatile method for simultaneously assessing the expression of a great number of RNA molecules in a dependable and reproducible manner (Goytain & Ng, 2020). This technology offers valuable advantages for ecological and evolutionary research, as it requires minimal RNA input and can handle lower-quality RNA, making it a more accessible option compared to RNA-seq. Importantly, it does not involve an amplification step and can detect even extremely low levels of RNA expression. In this study, an expanded Nanostring panel of probes was used, building upon a previous investigation (Kurko et al., 2020) that initially focused on gene expression related to age at maturity in Atlantic salmon. This updated panel includes > 100 additional genes, with a specific focus on a comprehensive collection of Hippo pathway components and other genes directly linked to the Hippo pathway. The selection process involved a combination of literature review and the use of tools like IPA (Ingenuity Pathway Analysis) from Qiagen, along with other web-based tools and databases (Kurko et al., 2020). Notably, the panel comprises probes targeting genes associated with age at maturity in Atlantic salmon, such as *vgll3a* on chromosome 25 and *six6a* on chromosome 9, along with their corresponding paralogs, *vgll3b* on chromosome 21, and *six6b* on chromosome 1. Additionally, the panel includes probes for other genes functionally connected to sexual maturation, including those within the HPG axis. Given that most candidate genes have multiple paralogs due to the duplicated Atlantic salmon genome (Lien et al., 2016), the paralogs of each gene of interest were integrated, identified through resources like SalmoBase (http://salmobase.org/) and the NCBI RefSeq databases. Further information regarding gene/paralog selection and nomenclature can be found in (Kurko et al., 2020). The gene accession numbers, symbols, full names, and functional classifications are provided in Supplementary file 1. The analysis of mRNA expression levels for these candidate genes involved the use of the Nanostring nCounter Analysis technology (NanoString Technologies, Seattle, WA, USA). Probes designed for each gene paralog, targeting all known transcript variants, were formulated using reference sequences from the NCBI RefSeq database. However, designing paralog-specific probes was not feasible for some genes due to high sequence similarity between paralogs. In practice, the execution involved the use of the nCounter Custom CodeSet for probes and the nCounter Master kit (NanoString Technologies). The RNA from each sample was denatured, followed by an overnight hybridization with the probes. Purification and image scanning were performed on the following day.

### Data analysis

For analyses of immature individuals, all nine candidate reference genes in the panel (*actb*, *ef1a* paralogs - *ef1aa*, *ef1ab*, and *ef1ac* – *gapdh*, *hprt1*, *prabc2* paralogs - *prabc2a* and *prabc2b*, and *rps20*) were used for data normalization due to their low coefficient of variation (CV) values across the samples. For analyses of mature individuals, one of the candidate reference genes, *actb*, was excluded because it displayed high variation (CV% > 50), and although it is commonly used as a reference gene in many studies, it appeared unsuitable for data normalization in the mature testis of Atlantic salmon. After this selection process, the raw count data obtained from Nanostring nCounter mRNA expression was normalized using RNA content normalization factors, which were calculated individually for each sample using the geometric mean count values of the chosen reference genes. Following normalization, a quality control assessment was conducted on the data, and all samples met the predefined threshold using the default criteria of nSolver Analysis Software v4.0 (NanoString Technologies; www.nanostring.com/products/nSolver). In the data analysis process using the software, the mean of the negative controls was subtracted, and positive control normalization was carried out by utilizing the geometric mean of all positive controls, following the manufacturer’s recommendations. To establish a baseline signal threshold, a normalized count value of 20 was set as the background signal. As a result, 114 genes in the panel displayed an average signal below this threshold across the samples, leaving 193 genes for subsequent analyses. Differential expression analysis was performed using the log-linear and negative binomial model (lm.nb function) integrated into Nanostring’s nSolver Advanced Analysis Module (nS/AAM). Predictive covariates inclu1ded the maturation status (*Immature 1*, *Immature 2*, *Mature*) and genotypes (*vgll3\**EE, *vgll3*\*LL), following nS/AAM’s recommendations. To address multiple hypothesis testing, the Benjamini-Yekutieli method (Benjamini & Yekutieli, 2001) was applied within the software, with an adjusted p-value threshold (FDR) of < 0.05 considered statistically significant (supplementary file 2). Additionally, for further exploration, log-transformed expression values were used to calculate pairwise Pearson correlation coefficients (r) between the gene expression variation of each candidate gene and GSI variation across all the samples.

We utilized the Weighted Gene Coexpression Network Analysis (WGCNA) R-package version 1.68 in R-package version 5.2.1 to identify gene co-expression networks (GCN) following the method by Langfelder & Horvath (2008) to compare the difference between the genotypes. To understand sample relationships, we hierarchically clustered samples based on gene expression. The process of building coexpression networks involved: (a) Calculating gene coexpressions using Pearson correlation coefficients, (b) creating an adjacency matrix to establish scale-free topology, (c) computing the topological overlap distance matrix based on the adjacency matrix, (d) hierarchical clustering of genes using the topological overlap distance, (e) identifying coexpressed gene modules using the cutTreeDynamic function with a minimum module size of 10 genes, (f) assigning colors to each module and representing module-specific expression profiles through the principal component (module eigengene), and finally (g) merging highly similar modules based on module eigengene (ME) dissimilarity with a distance threshold of 0.25 to finalize the coexpressed gene modules as described by Singh et al., 2021. Additionally, we conducted conditional coexpression analysis, constructing separate coexpression networks for each *vgll3* genotype to assess the preservation of modules between *vgll3*EEvgll3*LL* genotypes. We used a softpower of seven to create an adjacency matrix. To evaluate module preservation, we computed module preservation statistics using WGCNA, following the methodology of Langfelder, Luo, Oldham, & Horvath, 2011. A permutation test shuffled genes in the query network and calculated Z-scores. The individual Z-scores from 200 permutations were summarized into a Zsummary statistic.

To anticipate potential gene interactions and identify key genes with the highest interaction count, known as interacting hubs, we converted the differentially expressed genes in each comparison into their conserved orthologs in humans. We chose these orthologs due to their well-established and extensively studied interactome data across vertebrates. We then used these orthologs as input for STRING version 12.0, a comprehensive knowledge-based interactome database for vertebrates (Szklarczyk et al., 2023). Predicted gene interactions were based on various factors such as structural similarities, cellular co-localization, biochemical interactions, and co-regulation. We maintained a medium confidence level for each interaction or molecular connection prediction, which is the default threshold.

## Results

### Testicular gene expression differences between *vgll3* genotypes

Initially, we assessed variations in gene expression among the alternative *vgll3*EE* and *vgll3*LL* genotypes at the three distinct developmental stages described in the methods: *Immature 1, Immature 2* and *Mature* (as shown in Fig. 2). Between the genotypes differentially expressed (DE) genes were exhibited at *Immature 1*, with seven DE genes, and at *Immature 2*, with five DE genes, and 19 DE genes at the *Mature* stage. At *Immature 1*, all the DE genes, except *edar*, displayed higher expression in *vgll3*EE* genotype individuals (Fig. 2A). Our investigation into molecular interactions revealed that while only one gene (*tead3a*) had a direct interaction with *vgll3*, five genes exhibited potential direct interactions with *yap1* (*edar, pparac, rock2b, stk3a,* and *tead3a*) (as indicated by connecting lines in Fig. 2B). At *Immature 2*, we observed two genes (*admb* and *cebpba*) with higher expression in *vgll3*EE* genotype individuals, and three genes (*klf15d*, *limk2b*, and *rhoae*) showed increased expression in *vgll3*LL* genotype individuals (as illustrated in Fig. 2C). The predicted interactions indicated that one gene (*rhoae*) had a direct molecular interaction with *yap1*, while no gene exhibited direct interactions with *vgll3* (as shown in Fig. 2D). Moreover, the *rhoae* gene, which interacted with *yap1*, formed interacting hubs with two other DE genes (*limk2b* and *cebpba*) (Fig. 2D). At the *Mature* stage, we identified seven genes with higher expression in *vgll3*EE* genotype individuals and 12 genes that displayed increased expression in *vgll3*LL* genotype individuals (as presented in Fig. 2E). Our interaction analysis unveiled five genes (*kdm5bc*, *hif1a*, *mob1aa*, *rhoae*, and *six4*) with direct interactions with *yap1*, and two genes (*kdm5bc* and *akap11a*) with direct interactions with *vgll3* (as indicated in Fig. 2F). Among the genes with direct interactions with *yap1*, two genes, *rhoae* and *hif1a*, formed interacting hubs, suggesting multiple interactions with other DE genes (Fig. 2F). Ultimately, across all the comparisons, only one gene, *rhoae*, displayed differential expression in more than one comparison (at *Immature 2* and *Mature*). However, it exhibited lower expression in *vgll3*EE* at *Immature 2* but higher expression in *vgll3*EE* genotype individuals at the *Mature* stage.

**Figure 2.**
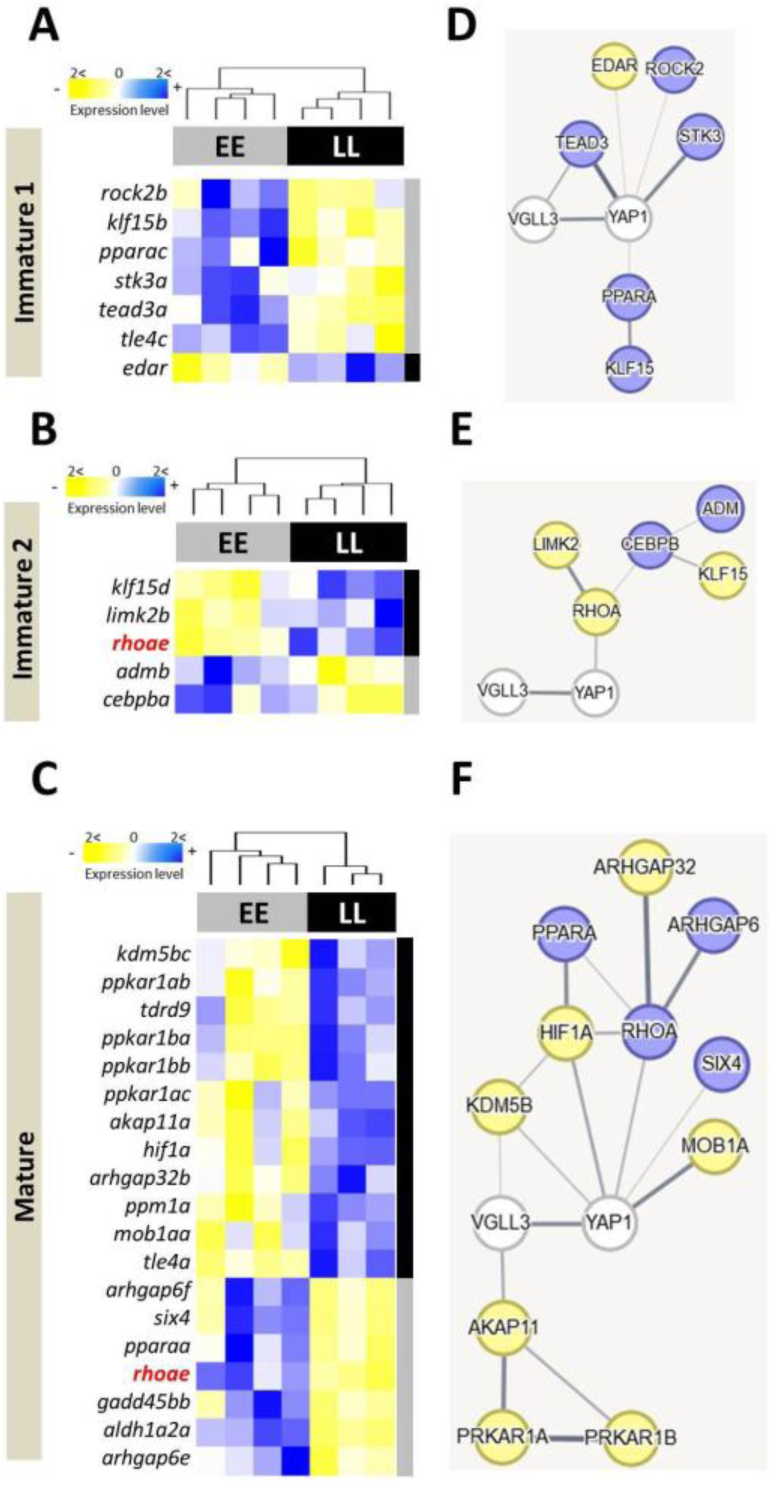
Testicular gene expression differences and predicted interactions between early and late maturation genotypes of *vgll3* in Atlantic salmon. Heatmaps illustrate genes with varying expression levels among alternative homozygous *vgll3* genotypes in Atlantic salmon testis at three developmental timepoints (**A-C**). Anticipated molecular connections between these genes are shown using STRING v12 (http://string-db.org/) (**D-F**). The thickness of the lines connecting the genes reflects the relative likelihood of interaction (thicker = more likely), while genes colored in blue and yellow in the predicted network signify higher and lower expression in *vgll3*EE* individuals, respectively.

### Testicular gene expression differences between maturation stages

To identify differences in gene expression between immature and mature testes, we compared immature individuals collected at both *Immature 1* and *Immature 2* stages with those collected at the *Mature* stage. We identified 185 DE genes between mature and immature individuals when both *vgll3* genotypes were combined. When analyzing each genotype separately, we found 154 DE genes in *vgll3*LL* genotype individuals and 173 DE genes in *vgll3*EE* genotype individuals (Fig. 3A). Overall, most DE genes in all three comparisons showed average lower expression at the *Mature* stage. In *vgll3*LL* individuals, only 23 of the 154 DE genes had higher expression in mature compared to immature individuals, while in *vgll3*EE* individuals, only 14 of the 173 DE genes showed higher average expression in the *Mature* stage individuals. Furthermore, 143 DE genes were found to overlap across all three comparisons, indicating differential expression independent of *vgll3* genotype (Fig. 3A). We observed more genotype-dependent DE genes in *vgll3*EE* individuals (30 genes, shown in red) than in *vgll3*LL* individuals (11 genes, shown in green in Fig. 3A). Importantly, the *vgll3a* paralog gene showed reduced expression at the *Mature* stage across all comparisons. However, *yap1*, the primary competitor of *vgll3* in the Hippo pathway, displayed reduced expression at the *Mature* stage only in *vgll3*EE*. These results suggest an overall reduction in expression of Hippo pathway components and interacting partners during testicular maturation, with minor differences between the *vgll3* genotypes.

**Figure 3.**
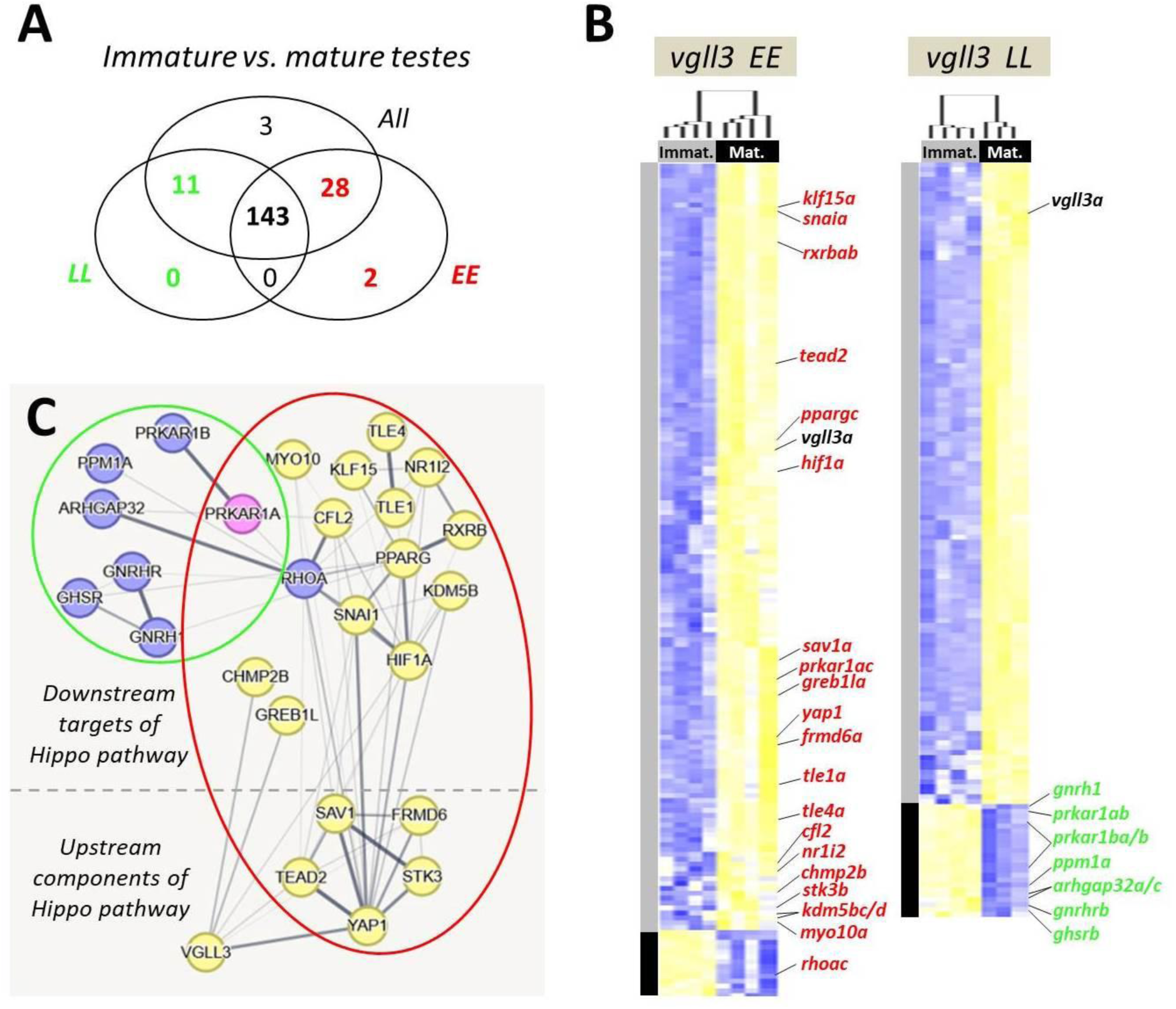
Testicular gene expression differences between immature versus mature Atlantic salmon. Venn diagram showing the numbers of differentially expressed genes overlapping between the comparisons (**A**). Heatmaps representing differentially expressed genes between the immature versus mature testes within alternative *vgll3* homozygotes (blue and yellow shadings represent higher and lower expressions, respectively) (**B**). Predicted interactions between the genes colored in green and red indicating the overlapping genes in the Venn diagram (**C**). The thickness of the connecting lines between the genes indicates the probability of the interaction (thicker = higher).

We proceeded to investigate potential functional/molecular interactions among DE genes between the *Immature* and *Mature* stages that exhibited *vgll3* genotype-specific variation in expression (colored numbers in Fig. 3A). The predicted interactions between these genes revealed that from the *vgll3* genotype specific comparisons, seven and 19 DE genes within *vgll3*LL* and *vgll3*EE* individuals, respectively, were found to be connected in an interaction network (colored green and red in Fig. 3B and C). Among the DE genes in *vgll3*LL* individuals, none had direct interactions with *vgll3* or *yap1*, and all showed higher expression at the *Immature* stage (Fig. 3E). Among the DE genes in *vgll3*EE* individuals, *greb1la*, *chmp2b*, *hif1a* and *tead2*, had direct interactions with *vgll3,* and all but three genes (*greb1la*, *chmp2b and klf15a*), had direct interactions with *yap1*. Moreover, all of these genes except *rhoac* showed reduced expression at the *Immature* stage contrary to the DE genes in *vgll3*LL*. Among the DE genes in *vgll3*LL* individuals none of them were upstream regulators of the Hippo pathway whereas in *vgll3*EE* individuals five key upstream regulators of the pathway were present (*frmd6a*, *sav1a, stk3b*, *tead2* and *yap1*). Finally, none of the DE genes in *vgll3*LL* individuals formed an interacting hub while several of hub genes were found among the DE genes in *vgll3*EE* such as *yap1, ppargc, rhoac, snaia* and *hif1a*. In general, these findings indicate more Hippo-dependent changes in the testis of mature vs. immature *vgll3*EE* individuals and a potential master regulatory role of *yap1* in this transition (Fig. 3C).

We observed paralog-specific expression patterns among the DE genes unique to both *vgll3* genotypes. For instance, while two paralogs of the *arhgap32* gene (*arhgap32d* and *arhgap32f*) showed increased expression at the *Mature* stage in both genotypes, two other paralogs, *arhgap32a* and *arhgap32c*, exhibited such increase only in the *vgll3*LL* genotype. This genotype-dependent divergence in paralog expression was also observed among other Hippo pathway interacting partners. However, most of these differences were present in *vgll3*EE* individuals, including *fgf14b* (vs. *fgf14a*), *frmd6a* (vs. *frmd6b/c*), *fstl3b* (vs. *fstl3a*), *greb1lb* (vs. *greb1la*), *kdm5ba/c/d* (vs. *kdm5bb*), *klf15a/d* (vs. *klf15b*), *myo10a* (vs. *myo10b/c/d/f*), *rhoac* (vs. *rhoaf/g/h*), *pcdh18a* (vs. *pcdh18b/c*), *rxrbaa* (vs. *rxrbab/c*), *sav1a* (vs. *sav1b*), *stk3b* (vs. *stk3a*), *tle1a* (vs. *tle1b*), *tle4a* (vs. *tle4b*), and *snaia* (vs. *snaib*). In all these cases, the paralogs outside the parentheses exhibited genotype-specific expression divergence in *vgll3*EE* individuals. The direction of divergence in *vgll3*EE* was often towards reduced expression at the *Mature* stage and similar to the direction of the other paralogs which showed the same patterns but in both genotypes. *Rhoac* and *sav1a*, were two exceptions among them, *rhoac* showed increased expression at the *Mature* stage of *vgll3*EE* individuals, contrary to its paralogs (*rhoaf/g/h*). An opposite expression pattern was observed for *sav1a*, which displayed reduced expression at the *Mature* stage of *vgll3*\*EE individuals, in contrast to *sav1b*, which had increased expression at the *Mature* stage in both genotypes. A more extreme example was paralogous expression divergence of the *prkar1a* gene: *prkar1ab* had higher expression only in *vgll3*\*LL, while the other paralog, *prkar1ac*, showed lower expression only in *vgll3*EE* at the *Mature* stage. These observations suggest *vgll3* genotype-specific expression divergence in paralogs of the Hippo pathway-related genes during testicular maturation. In some cases, such as *prkar1a*, *rhoa*, and *sav1*, more pronounced divergence between genotypes may even indicate neo-functionalization between their paralogs.

### Genotype specific gene coexpression networks

To gain a better understanding of the transcriptional dynamics of Hippo pathway components and their interacting genes, we used network-based co-expression analyses. This approach enabled us to examine genotype-specific changes in each network. We first constructed gene co-expression networks (GCNs) for both *vgll3* genotypes in the testis. Next, we evaluated the conservation of gene co-expression modules between genotypes by building the GCN for one genotype (*vgll3*EE* or *vgll3*LL*) and testing module preservation in the other (*vgll3*LL* or *vgll3*EE*).

We found seven modules in *vgll3*EE* (Fig. 4A and B) of which two modules, green and red, showed low preservation (Zsummary < 1) in *vgll3*LL*, *i.e.* many of the genes in each module no longer had significant expression correlations in *vgll3*LL* genotype (the genes lacking color in Fig. 4C). In the red module, eight out of 16 genes showed no coexpression preservation in *vgll3*LL* genotype individuals (Fig. 4C). Also in the green module, five out of 25 genes exhibited no coexpression preservation in *vgll3*LL* individuals (Fig. 4C). Using knowledge-based interactome prediction, we used the genes within each module to identify potential interactions among them. In both modules at least one gene, among those lacking coexpression preservation, had a direct interaction with *vgll3* (i.e., *greb1la* in the green and *six6a* in the red module) (Fig. 4C). Interestingly, in each module a different paralog of *vgll3* (*vgll3b* in the red and *vgll3a* in the green module) was found, however, both paralogs retained their coexpression preservation in *vgll3*LL* genotype individuals (Fig. 4C). Moreover, in each module one upstream Hippo component encoding gene with direct interaction with *yap1* also lacked coexpression preservation in *vgll3*LL* genotype individuals (*lats1b* in the green and *sav1a* in the red module) (Fig. 4C).

**Figure 4.**
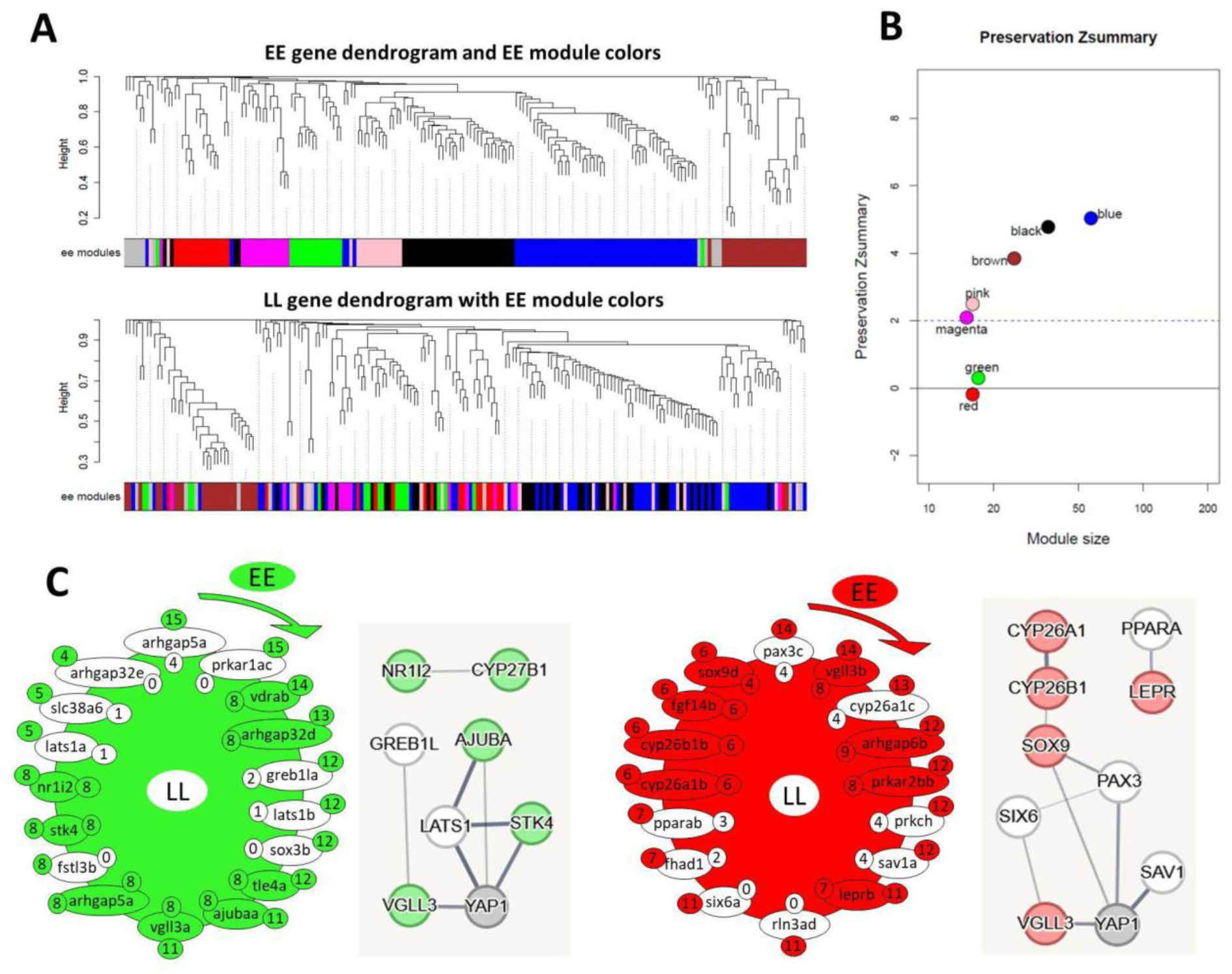
Coexpression analysis of *vgll3*EE* genotype in salmon testis. Dendrograms display clustering and preservation of *vgll3*EE* modules in *vgll3*LL* individuals, highlighting the lack of preservation in the top panels (**A**). Zsummary scores indicate low preservation (<2) and moderate preservation (2 to 6) for *vgll3*EE* modules in *vgll3*LL* (**B**). The genes in the least preserved *vgll3*EE* module in *vgll3*LL* individuals (**C**). Non-preserved genes are shown without color, and clockwise arrows indicate directions from the most to the least connected genes within the module, the numbers beside the genes and outside the module represent the number of expression correlations for each gene (with other genes within the same module) in *vgll3*EE* modules and numbers inside represent the number of expression correlations for each gene preserved in *vgll3*LL*. Predicted interactions between the genes within each of the least preserved modules are depicted on the right side of the modules. Line thickness is relative to interaction probabilities (thicker = higher).

In *vgll3*LL* compared to *vgll3*EE* individuals, we identified five modules, two of which (red and brown) showed low preservation levels (Zsummary < 1) (Fig. 5A, B). Within the red module, interaction predictions revealed two genes, *ets1b* and *greb1la*, that directly interact with *vgll3*. Both genes lost their co-expression preservation in *vgll3*EE* individuals (Fig. 5C). Interestingly, *greb1la* was also the most connected gene in this module, showing significant expression correlation with 19 out of 20 genes. Moreover, *stk3b*, an upstream regulator of the Hippo pathway with a direct interaction with *yap1*, also lacked expression correlation in *vgll3*EE* individuals. In the brown module, three genes, *six6a*, *akap11b*, and *kdm5bd*, had direct interactions with *vgll3* and exhibited a loss of co-expression with other module genes in *vgll3*EE* individuals (Fig. 5C). Similarly, three upstream regulators of the Hippo pathway, *lats1a*, *lats1b*, and *sav1b*, which directly interact with *yap1*, also lacked expression correlation in *vgll3*EE* individuals.

**Figure 5.**
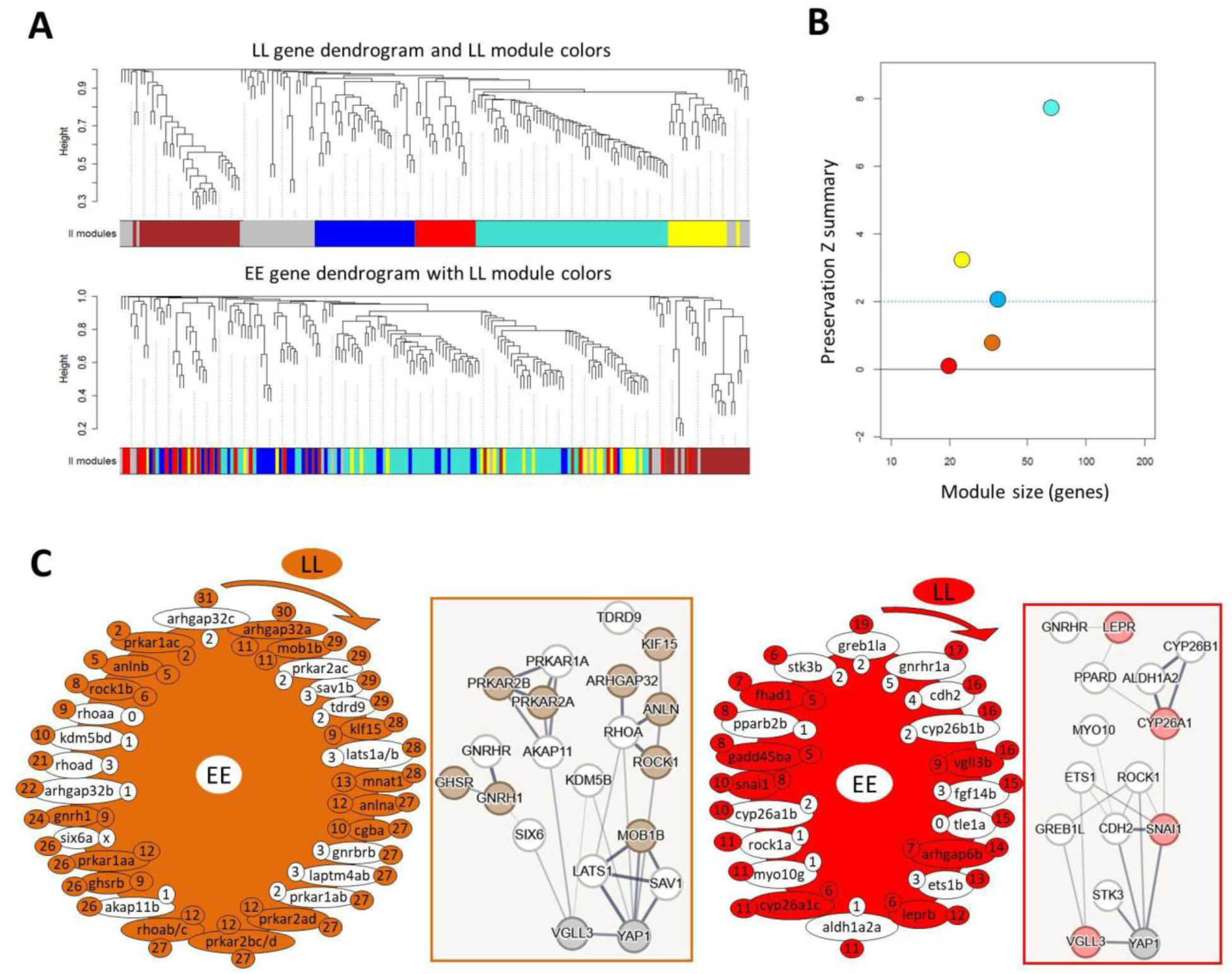
Coexpression analysis of *vgll3*LL* genotype in salmon testis. Dendrograms display clustering and preservation of *vgll3*LL* modules in *vgll3*EE* individuals, highlighting the lack of preservation in the top panels (**A**). Zsummary scores indicate low preservation (<2) and moderate preservation (2 to 6) for *vgll3*LL* modules in *vgll3*EE* (B). The genes in the least preserved *vgll3*LL* module in *vgll3*EE* (**C**). Non-preserved genes are shown without color, and clockwise arrows indicate directions from the most to the least connected genes within the module, the numbers besides the genes and outside the module represent number of expression correlation for each gene (with other genes within the same module) in *vgll3*LL* modules and numbers inside represent the number of expression correlations for each gene preserved in *vgll3*EE*. Predicted interactions between the genes within each of the least preserved modules are depicted on the right side of the modules. Line thickness is relative to interaction probabilities.

## Discussion

Sexual maturation, or puberty, is initiated in the brain, primarily when the hypothalamus secretes gonadotropin-releasing hormone (GnRH), which triggers the pituitary gland to release luteinizing hormone (LH) and follicle-stimulating hormone (FSH) (Herbison, 2016). The activation of the FSH signal initiates a cascade of signaling pathways that regulate various metabolic, developmental, and cellular processes essential for spermatogenesis (Ni et al., 2019, 2020). Among these pathways, the Hippo pathway has recently been identified as a signal directly regulated by FSH, playing significant roles not only in the early stages of testicular maturation but also in later stages of spermatogenesis and related pathologies (Abou Nader et al., 2019; Sen Sharma et al., 2019; Zhang et al., 2019; Meng et al., 2020; Yu et al., 2021; Onel et al., 2023). However, compared to other well-known pathways involved in this process, such as the cAMP-PKA, ERK1/2-MAPK, PI3K-AKT, and TGF-β signaling pathways (Ni et al., 2019, 2020), much less is understood about the specific molecular players and their functions within the Hippo pathway cascade. In this study, we explored how different genotypes of the *vgll3* gene impact transcriptional activity of the Hippo pathway components in immature and mature testis. We also examined whether *vgll3*-associated transcriptional changes in the testis could serve as early indicators of maturation, even before any visible signs of maturation become apparent in the testis. The molecular signaling regulated by the Hippo pathway is known to play essential and conserved roles from early stages of testicular development to final stages of spermatogenesis (Sun et al., 2008; Abou Nader et al., 2019; Sen Sharma et al., 2019; Onel et al., 2023). This allowed us to gain insights into how Hippo pathway signaling is regulated in the testes of Atlantic salmon, which have a reproductive strategy closely linked to seasonal variations and body fat reserves.

### Potential heterochrony in the Hippo-dependent regulation of early signals of spermatogenesis

The expression patterns of Hippo pathway components and their interacting partners across the three time points suggest a potential heterochronic shift in the timing of testicular development and spermatogenesis between the *vgll3* genotypes, with *vgll3*EE* individuals displaying more advanced patterns than *vgll3*LL* individuals. For example, at the earliest immature time-point (spring: *Immature-1*), we found the expression of genes encoding Hippo components/interacting partners with an enhancing role in early stages of spermatogenesis to be induced in the testis of *vgll3*EE* individuals. Notably, we found that *stk3a*, a paralog of a gene encoding a key kinase in the Hippo pathway (also known as *MST2*), is more highly expressed in the testis of individuals with the *vgll3*EE* than the *vgll3*LL* genotype at the earliest *Immature* stage (spring). In mice, MST2 activation is required for YAP suppression, which promotes the differentiation of the epididymal initial segment (IS) in the testis (Meng et al., 2020). The differentiation of the epididymal IS is considered an early sign of developmental changes in the testis, leading to the onset of testicular maturation and the increase in androgen (testosterone) levels (O’Hara et al., 2011). Consistently, we also found induction of a *KLF15* paralog gene (*klf15b*) in the *vgll3*EE* genotype, which is known as a direct target of androgen in Sertoli cells in mice (de Gendt et al., 2014). Other important genes encoding interacting partners of the Hippo pathway and displaying higher expression in the testes of individuals with the *vgll3*EE* genotype were *rock2b* and *pparac*. The first gene, *rock2b*, encodes a paralog of *ROCK* genes (that are essential for the activation of the ROCK/LIMK signaling pathway), which is required for the formation of cell-cell junctions between Sertoli cells and spermatogonia at the early stages of spermatogenesis (Lui et al., 2003a). The latter, *pparac*, is a paralog of a gene encoding a nuclear receptor that senses lipid availability in cells and plays a key role in lipid metabolism, ensuring sufficient energy for testis development, maturation, and spermatogenesis (Monrose et al., 2021). It has been demonstrated in mammalian studies that FSH secretion can upregulate the expression of both *MST2* and *PPARα* in the testis (Schultz et al., 1999; Sen Sharma et al., 2019). Previously, we also found early induction of *fshb* expression in the pituitary of male salmon with the *vgll3*EE* genotype (as early as one year old), long before any visible signs of maturation appeared in the testis (Ahi et al., 2022, 2023). This could explain the early induction of these genes (*stk3a/MST2* and *pparac*) in the testis during the springtime. However, the increased expression of *pparac* in the *vgll3*EE* genotype may partly be due to the higher availability of lipids in males with this genotype, as *PPARα* can be directly induced by circulating lipids (Hasan et al., 2020). This scenario also aligns with our previous findings in adipose tissue, which suggested a potentially greater lipid storage capacity in males with the *vgll3*EE* genotype at the same time points in spring (Ahi et al., 2024).

The observation that components of similar molecular signals involved in the early stages of testicular development and spermatogenesis showed increased expression in the vgll3*LL genotype at the later immature time point suggests a potential heterochrony between the *vgll3*EE* and *vgll3*LL* genotypes in the timing of these developmental processes. For example, we observed increased expression of components of the ROCK/LIMK signaling pathway, specifically *limk2b* and *rhoae*, in the testes of individuals with the *vgll3*LL* genotype. Both genes are involved in the formation of cell-cell junctions between Sertoli cells and spermatogonia at the early stages of spermatogenesis (Takahashi et al., 2002; Lui et al., 2003b, 2003a). Interestingly, another paralog of the *KLF15* gene (*klf15d*) was found to have higher expression in the *vgll3*LL* genotype at this time-point, suggesting a subsequent heterochronic shift in androgen effects on Sertoli cells in this genotype (de Gendt et al., 2014). However, more developmental time-points are needed to obtain a clearer and more detailed understanding of these molecular cascades during testicular maturation in both genotypes.

At the mature time point, heterochrony appears to be maintained, with genes involved in various stages of testicular development and spermatogenesis showing shifts, as suggested by early-stage genes being more highly expressed in *vgll3*LL* individuals, while later-stage markers have increased expression in *vgll3*EE* individuals. For instance, *kdm5b* (also known as *PLU-1* or *JARI1B*), a gene with essential role in pre-meiotic spermatogonial proliferation and repression of terminal spermatocyte differentiation (Simpson et al., 2005), had higher expression in the testis of *vgll3*LL* individuals. Other genes with essential role in earlier stages of spermatogenesis that had higher expression in *vgll3*LL* individuals include *tdrd9* and *ppm1a* (or *PP2AC*); both required for the initiation of meiotic phase (Shoji et al., 2009; Chen et al., 2022), and *prkar1* with critical functions during the early prophase I stage of meiosis (the pachytene stage) (Burton et al., 2006). In contrast, genes involved in the later stages of spermatogenesis had higher expression in the mature testis of *vgll3*EE* individuals such as *arhgap6*; required for actin remodeling and sperm motility (Almog et al., 2008), *aldh1a2*; highly expressed during spermatocyte differentiation and spermatid maturation (Griswold et al., 2022), and *ppara*; required for lipid metabolism and providing energy for sperm motility (Froment et al., 2006).

### Paralog-specific expression patterns during spermatogenesis

A paralog of an early meiosis stage marker, *prkar1ab*, exhibited higher expression in *vgll3*LL*, while the other paralog, *prkar1ac*, showed lower expression in *vgll3*EE* during the *Mature* stage. This indicates that the testis of *vgll3*LL* individuals is more active at earlier stages of spermatogenesis (Burton et al., 2006). Such paralog specificity followed the same trend of heterochrony, i.e., earlier-stage genes having higher expression in *vgll3*LL* at each time point (and vice versa).For instance, we observed the reduced expression of genes known to be more active in earlier stages of spermatogenesis in the testis of *vgll3*EE* individuals during the *Mature* stage. Among these were: three *kdm5b* paralogs (*kdm5ba/c/d*), which act as inducers of spermatogonial proliferation and repressors of terminal spermatocyte differentiation (Simpson et al., 2005); two *klf15* paralogs (*klf15a/d*), markers of early-stage androgen sensing in Sertoli cells (de Gendt et al., 2014); *myo10a* paralog, a marker of mid-spermatogenesis stages and spindle formation during meiosis (Brieño-Enríquez et al., 2017); *rhoac* paralog, a marker of cell junctions in spermatogonia (Lui et al., 2003b, 2003a); *stk3b* paralog, a marker of earlier stages of testicular development (Meng et al., 2020); *pcdh18a* paralog, a marker of the pre-meiotic stage of spermatogenesis (Jan et al., 2017); *tle4a* and *snaia* paralogs, markers of spermatogonial differentiation (Micati et al., 2018; Zhang et al., 2021). These paralog-specific patterns align with earlier findings, which showed that markers of earlier stages of spermatogenesis are more active in the testis of *vgll3*LL* individuals. Moreover, these expression differences between the paralogs suggest potential sub-functionalization among these paralogs, which appears to be influenced by distinct Hippo pathway activity.

### Different Hippo pathway components and interacting partners may participate in the testicular maturation shift

By combining co-expression analysis and interactome prediction, we identified co-expression modules most affected by differences between the *vgll3* genotypes. This approach also allowed us to pinpoint potential key contributors to the distinct gene expression patterns observed between the genotypes. Importantly, within the most affected gene co-expression networks (the least preserved), we identified distinct *vgll3* paralogs. While the co-expression of these paralogs remained unaffected, several key interacting partners and Hippo pathway components appeared to be differentially affected between genotypes. Among the affected upstream components of the Hippo pathway were *lats1*, *sav1a*, and *stk3b*, which have been previously discussed for their roles in testicular development and spermatogenesis (see above). Downstream effectors of the Hippo pathway showing direct connections with *vgll3* included *kdm5bd* (as described above) and *ets1b*. Recent research indicates that Vgll3 can directly bind to regulatory sequences of the *ets1* gene in salmon testes (Verta et al., 2024). ETS1, a transcription factor, has been implicated in mammalian spermatogonia stem cell differentiation (Lu et al., 2024) and is also present in mature spermatozoa (Pittoggi et al., 2001). Furthermore, ETS1 binding sites have been identified in enhancers driving mitotic (but not meiotic) spermatogenesis in mammals (Maezawa et al., 2020). In our study, we found a genotype-specific expression correlation between *ets1b* and *vgll3b* paralog, observed only in the *vgll3*LL* genotype. This may suggest differences in the initiation of spermatogonial proliferation and differentiation between genotypes. Moreover, we identified three genes, *greb1la*, *six6a*, and *akap11b*, that directly interact with *vgll3* but are not classified as Hippo pathway components. These genes are implicated in life-history traits in salmonids. For instance, *greb1l* has been associated with migration timing in Chinook salmon and Steelhead (Collins et al., 2020; Koch & Narum, 2020; Horn & Narum, 2023), while *six6* and *akap11* have been linked to age-at-maturity in Atlantic salmon (Barson et al., 2015; Kurko et al., 2020; Moustakas-Verho et al., 2020; Sinclair-Waters et al., 2020). In mice, the antisense transcript of *SIX6* (*SIX6OS1*) is required for testicular development and spermatogenesis (Gómez-H et al., 2016), but the specific role of *SIX6* in testicular function remains unexplored in vertebrates. Similarly, *GREB1L* has recently been proposed as a marker for male infertility in mammals (Lillepea et al., 2024), and its paralog *GREB1* is involved in estrogen-dependent functions of Sertoli cells (Lin et al., 2014). However, the precise role of *greb1l* in vertebrate spermatogenesis remains unclear. Lastly, *AKAP11* (also known as *hAKAP220*) is expressed throughout spermatogenesis and is believed to play a role in cytoskeletal structure formation (Reinton et al., 2000). These findings emphasize the significance of *vgll3* in testicular maturation, extending beyond its conventional role as a co-factor of the Hippo pathway. As suggested, *vgll3* may influence testicular processes by interacting with other signaling pathways (Verta et al., 2024).

## Conclusions

This study offers molecular evidence that connects testicular gene expression with genotype-related variation in life-history strategies, particularly related to sexual maturation. Our results provide evidence of gene expression that supports the ability to predict early maturation in the testis before visible changes linked to maturation manifest in immature male Atlantic salmon. This predictability is influenced by a unique transcriptional signature involving components and interacting partners of the Hippo pathway, which appears significantly affected in individuals with both *early* and *late vgll3* genotypes. Moreover, at each compared stage, the testis of individuals with *early vgll3* genotype seems to have a transcriptional signature suggestive of more advanced stages of spermatogenesis, indicating potential heterochronic shift. These findings may imply that particular elements within the Hippo signaling pathway as well as less known factors with potential interactions with the pathway (e.g., *greb1lb* and *six6a*) could play a substantial role in driving the varying effects of *vgll3* genotypes on testicular gene expression and the timing of maturation. Moreover, given that the Hippo pathway is an environmental sensing pathway, particularly responsive to food availability and temperature, both of which influence maturation, these findings suggest a likely link between the environment and heterochronic events through the *vgll3*/Hippo axis, leading to differences in age at maturity. However, further functional assessments are needed to confirm these proposed distinctions.

## Acknowledgements

We thank Jaakko Erkinaro and Natural Resources Institute, Finland (Luke) for providing access to mature salmon used for creating the families used in this study, and Pooja Singh for guiding us through the WGCNA data analysis. We also thank N. Piavchenko, S. Andrew, O. Andersson, T. Aykanat, Y. Czorlich, A. House, M. Lindqvist, N. Lorenzen, O. Mehtälä, K. Mobley, J. Moustakas-Verho, O. Ovaskainen, S. Papakostas, N. Parre, A. Ruokolainen, V. Pritchard, K. Salminen, M. Sinclair-Waters, S. Tillanen, and K. Zueva for help related to gamete stripping, sample processing, tagging, genotyping, phenotyping or fish husbandry, and the staff at the Natural Resources Institute Finland (Luke) hatchery in Taivalkoski hatchery for help during spawning.

## Author Contributions

EPA, CRP, JPV, JK and PVD conceived the study; JPV, PVD and CRP reared and sampled the fish; CRP provided resources; AR, EPA and JPV performed experiments; EPA, JPV and JK developed methodology and analyzed the data; EPA, JPV and CRP interpreted results of the experiments; EPA, JPV and CRP drafted the manuscript, with EPA having the main contribution, and all authors approved the final version of manuscript.

## Ethical approval

Animal experimentation followed European Union Directive 2010/63/EU under license ESAVI/35841/2020 granted by the Animal Experiment Board in Finland (ELLA).

## Data availability

All the gene expression data generated during this study are included in this article as supplementary file.

**Supplementary Data: Expression data and statistical analysis.**

